# Short-Term Plasticity Following Motor Sequence Learning Revealed by Diffusion MRI

**DOI:** 10.1101/553628

**Authors:** Ido Tavor, Rotem Botvinik-Nezer, Michal Bernstein-Eliav, Galia Tsarfaty, Yaniv Assaf

## Abstract

Current non-invasive methods to detect structural plasticity in humans are mainly used to study long-term changes. Diffusion MRI was recently proposed as a novel approach to reveal gray matter changes following spatial navigation learning and object-location memory tasks. In the present work we used diffusion MRI to investigate the short-term neuroplasticity that accompanies sequential motor learning. Following a 45-minutes training session in which participants learned to accurately play a short sequence on a piano keyboard, changes in diffusion properties were revealed mainly in motor system regions such as the premotor cortex and cerebellum. A second learning session taking place immediately afterwards shifted the attention of participants onto timing of key pressing instead of accuracy. This second session induced a different plasticity pattern, demonstrating the dynamic nature of this phenomenon, formerly thought to require months of training in order to be detectable. These results provide us with an important reminder that the brain is an extremely dynamic structure. Furthermore, diffusion MRI offers a novel measure to follow tissue plasticity particularly over short timescales, allowing new insights into the dynamics of structural brain plasticity.

## 1. Introduction

Neuroplasticity, the ability of the nervous system to adapt its organization according to the dynamic internal and external environment, has been extensively studied in recent decades. Numerous experiments have demonstrated neural plasticity throughout the brain, both functionally and structurally. However, structural plasticity was mainly investigated over long timescales such as months or weeks, using conventional anatomical magnetic resonance imaging (MRI) (Draganski et al., 2004; Scholz et al., 2009; Zatorre et al., 2012).

Recently, diffusion-weighted (DW) MRI provided a new approach to explore short-term neuroplasticity in the human brain (Assaf, 2018; Brodt et al., 2018; Blumenfeld-Katzir et al., 2011; Hofstetter et al., 2013, 2017; Sagi et al., 2012; Scholz et al., 2009; Tavor et al., 2013). The mean diffusivity (MD) of water molecules, extracted from DW-MRI, has been shown to serve as a highly-sensitive biomarker for microstructural changes associated with several types of learning: spatial navigation (Sagi et al., 2012; Tavor et al., 2013), phonological language learning (Hofstetter et al., 2017), and object-location memory (Brodt et al., 2018). Structural modifications were measured by a decrease in gray matter's mean diffusivity (MD) and were detectable within hours of learning in brain regions that were relevant to the investigated cognitive domain (e.g. spatial navigation induced MD decrease in the hippocampus, reflecting microstructural changes). In the present study we employed this novel approach to study short-term structural plasticity following a complex, multi-step motor learning task.

The motor system has been the subject of many neural plasticity studies in recent years (Bezzola et al., 2011; Doyon and Benali, 2005; Draganski et al., 2004; Herholz et al., 2016; Muellbacher et al., 2001; Sale et al., 2017; Sanes and Donoghue, 2000; Scholz et al., 2009). Using various imaging techniques, it is possible to follow on both structural (Bezzola et al., 2011; Draganski et al., 2004; Rudebeck et al., 2009) and functional (Poldrack, 2000; Reithler et al., 2010; Ungerleider et al., 2002) brain changes, mainly as a result of learning and memory of motor-related procedures. When learning a new motor skill, improvement is first seen in accuracy, and after extensive training also in the speed and rhythm of performance (Doyon, 2008; Hikosaka et al., 2002; Lehéricy et al., 2005). Different parts of the motor cortex were found to be activated in early stages of learning as opposed to later stages (Hikosaka et al., 2002; Lehéricy et al., 2005). This dissociation exists also in the basal nuclei, in which the anterior striatum was found to be involved in the acquisition of new motor skills, whereas the posterior striatum may be critical for the long-term storage of those skilled behaviors (Lehéricy et al., 2005). The cerebellum and its connections to the cortex were shown to contribute to different aspects of motor learning as well (Doyon et al., 2003).

While a variety of studies focused on the functional aspects of motor plasticity, little is known about the structural remodeling of the tissue, particularly in the short-term. In this study we used diffusion MRI to investigate the short-term neuroplasticity that accompanies sequential motor learning. Specifically, we set up a motor-sequence learning task using an electric piano keyboard. Thirty-two non-musician participants were scanned before and immediately after a 45-minutes training session in which they learned to play a short sequence based on Beethoven's Für Elise. Participants were instructed to repeat the sequence with an increasing number of notes and were given feedback on the accuracy of their key pressing. A subset of fifteen participants continued on to a second learning session in which they received feedback on the rhythm, rather than accuracy, of key pressing (this second stage is only possible after reaching a sufficiently high accuracy level). Finally, these fifteen participants were scanned for the third time. To control for the localization of brain changes we also acquired a preliminary cohort of 8 professional pianists that were scanned before and after performing the same task.

We hypothesized that a short-term motor-sequence learning task will induce structural brain changes reflected as decreased mean diffusivity within several motor system regions. Furthermore, we hypothesized that a second learning session, which emphasizes different aspects of motor learning (i.e. rhythm as opposed to accuracy), will result in different patterns of structural plasticity, and that the professional pianists’ cohort will exhibit yet another pattern of brain changes associated with the same piano-learning task.

## 2. Material and Methods

### 2.1. Participants

Forty healthy volunteers participated in this study (mean age 25.7; S.D. 3.1, 20 males, all right-handed), with no history of neurological disease, psychological disorders, drug or alcohol abuse, or use of neuropsychiatric medication. The research protocol was approved by the Institutional Review Board of the Sourasky and Sheba Medical Centers. All participants signed an informed consent form. Out of the whole cohort, 8 participants were professional pianists (with formal musical education and more than 8 years of experience). Out of the 32 naïve participants, 17 performed a single learning session, and 15 continued to a second learning session.

### 2.2. Learning Task

During the task participants learned to play a short sequence on an electric piano keyboard (MEDELI M10, Medeli Electronics Co.). The training sequence was the first 51 notes of the right-hand part of Beethoven's *Für Elise*. Using an in-house software, participants were presented with an increasing number of notes (from 1 to 51) on a virtual keyboard and were instructed to repeat the sequence on the keyboard in front of them using their right hand. The presentation of a sequence included both visual and auditory stimuli. After repeating the sequence participants were given feedback on their accuracy. The learning session included 63 trials and lasted approximately 45 minutes.

A subset of 15 participants continued to a second learning session, in which they played the entire 51-notes sequence over and over again and were given feedback on the rhythm of key pressing, rather than accuracy. A key press was considered a rhythm error when it differed in time from the correct piece in more than an eighth of a beat. The second learning session included 25 trials and lasted approximately 40 minutes.

### 2.3. MRI Acquisition

MRI was performed using a GE Signa 3.0T scanner (GE, Milwaukee, USA). Participants underwent two or three MRI scans, before and immediately after each learning session (see Section 2.2 above). Thus, the scans were approximately an hour apart. The MRI protocol of each of the two scans included DTI and conventional anatomical sequences for radiological screening, all acquired with an 8-channel head-coil.

#### 2.3.1. Conventional Anatomical Sequences

T1-weighted images were acquired with a 3D spoiled gradient-recalled echo (SPGR) sequence with the following parameters: up to 160 axial slices (whole-brain coverage), TR/TE = 9/3 ms, resolution 1 × 1 × 1 mm^3^, scan time 4 minutes. In addition to the T1-weighted scan, T2-weighted images (TR/TE = 6,500/85) and FLAIR images (TR/TE/TI = 9,000/140/2,100) were acquired. The entire anatomical data set was used for radiological screening.

#### 2.3.2. DTI Protocol

Spin-echo diffusion weighted echo-planar imaging sequences were performed with up to 70 axial slices (to cover the whole brain) and resolution of 2.1 × 2.1 × 2.1 mm^3^ reconstructed to 1.58 × 1.58 × 2.1 mm^3^ (field of view is 202 mm^2^ and acquisition matrix dimension is 96 × 96 reconstructed to 128 × 128). Diffusion parameters were: ∆/δ= 33/26 ms, b value of 1000 s/mm^2^ was acquired with 30 gradient directions and an additional image is obtained with no diffusion weighting (b_0_ image).

### 2.4. Behavioral Data Analysis

For each trial, two measurements were calculated to distinguish between two different learning aspects: (1) accuracy of key pressing was measured by the number of correct notes that were played in each trial and (2) accuracy of rhythm was measured by the average error in time (the distance between the original timing of each note and the time pressed by the participant) per note for each trial. It is noted that the first measurement is limited to the number of notes that are included in each trial. The second measurement is also influenced by the number of notes to be played in each trial (as the error in time is accumulating throughout the musical part). To prevent this from influencing our measure, timing was measured relatively to the previous note and averaged across all notes in a given trial.

### 2.5. MRI Data Analysis

DW images were corrected for motion using a least-squares algorithm and 6-parameter (rigid-body) transformations implemented in SPM software. Then, DTI analysis was performed using an in-house software implemented in MATLAB 8.4.0 (<u>Mathworks, Natick, MA</u>), from which maps of mean diffusivity (MD) were computed.

### 2.6. Image Processing

Image processing including registration, spatial normalization and spatial smoothing were done using SPM as described previously (Sagi et al., 2012; Tavor et al., 2013). Briefly, to ensure optimal alignment for voxel-based statistics we used the fractional anisotropy (FA) maps in a two-step spatial registration routine. Non-linear transformations were used for both within-subject and between-subject registration (See Sagi et al. 2012 for more details).

### 2.7. Statistical Analysis

Voxel-based statistics is used to detect regionally specific differences in brain tissue on a voxel-by-voxel basis. Statistical voxel-based analysis of MD maps was performed using MATLAB. Specifically, MD maps acquired before and after the learning task were compared in order to detect brain regions in which the mean diffusivity decreased following the task, indicating a structural change. The decrease in mean diffusivity was considered statistically significant in clusters that exceeded a threshold of *P* < 0.005 uncorrected and cluster size of 37 voxels, which is equivalent to a corrected threshold of *P* < 0.05 according to Monte Carlo simulation implemented in 3dClustSim program in the AFNI software package (i.e. the probability to find false positive clusters in this size is less than 5%; see *technical considerations* in the discussion section).

## 3. Results

### 3.1. Behavioral Results

#### 3.1.1. First Learning Session

All participants improved their accuracy from trial to trial and by the end of the first learning session succeeded to play most of the notes correctly. On average, participants played 46.6 ± 1.49 (mean ± S.E.M) correct notes out of the total 51 notes at the end of the task (Figure 1). While improvement in accuracy was relatively high, participants' timing barely improved during the first learning session: as the number of notes they had to play increased, the task became more difficult in the aspect of timing and the error in time increased. Starting from trial 16, in which participants had to play 13 notes, the average error in time was approximately 150 milliseconds per note. Participants did not improve in this aspect during the rest of the task, even in trials in which the number of notes was fixed (Figure 2A). In line with previous studies, improvement in accuracy preceded improvement in timing in our task, which was probably too short in order to reach advanced stages of motor skill learning. It should be noted that during the first learning session participants were only instructed to play the correct notes and were not given feedback on their timing.

**Figure 1:**
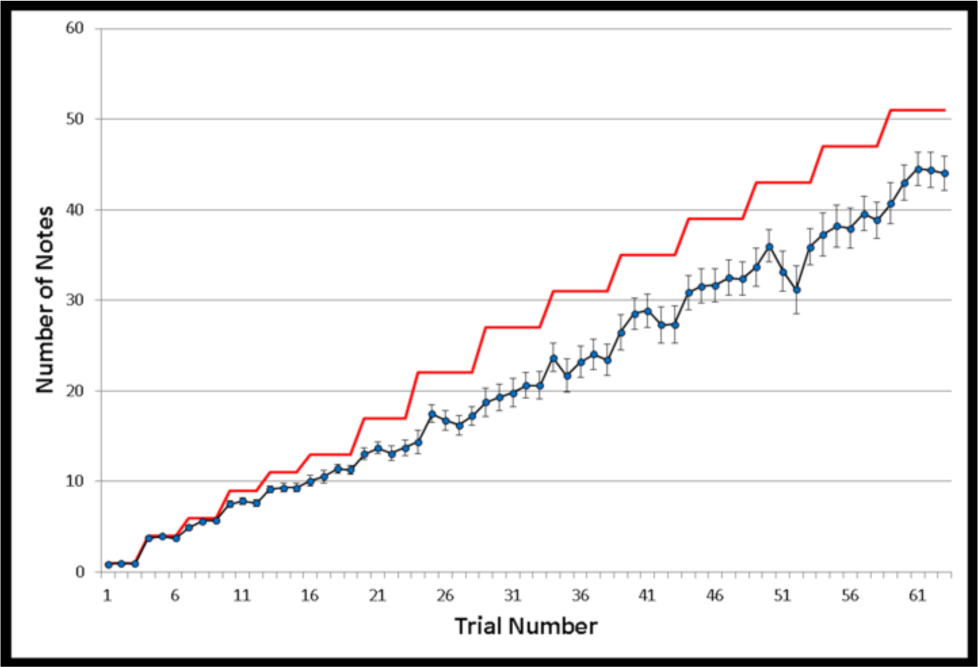
Performance in Piano Training (Accuracy) Accuracy of key pressing during the first learning session is shown. Blue circles represent the average number of correct notes played by participants in each trial. The number of notes that were presented to participants in each trial is shown in red. On average, the best performance participants achieved was 46.6 correct notes out of 51. Error bars depict standard error of mean.

**Figure 2:**
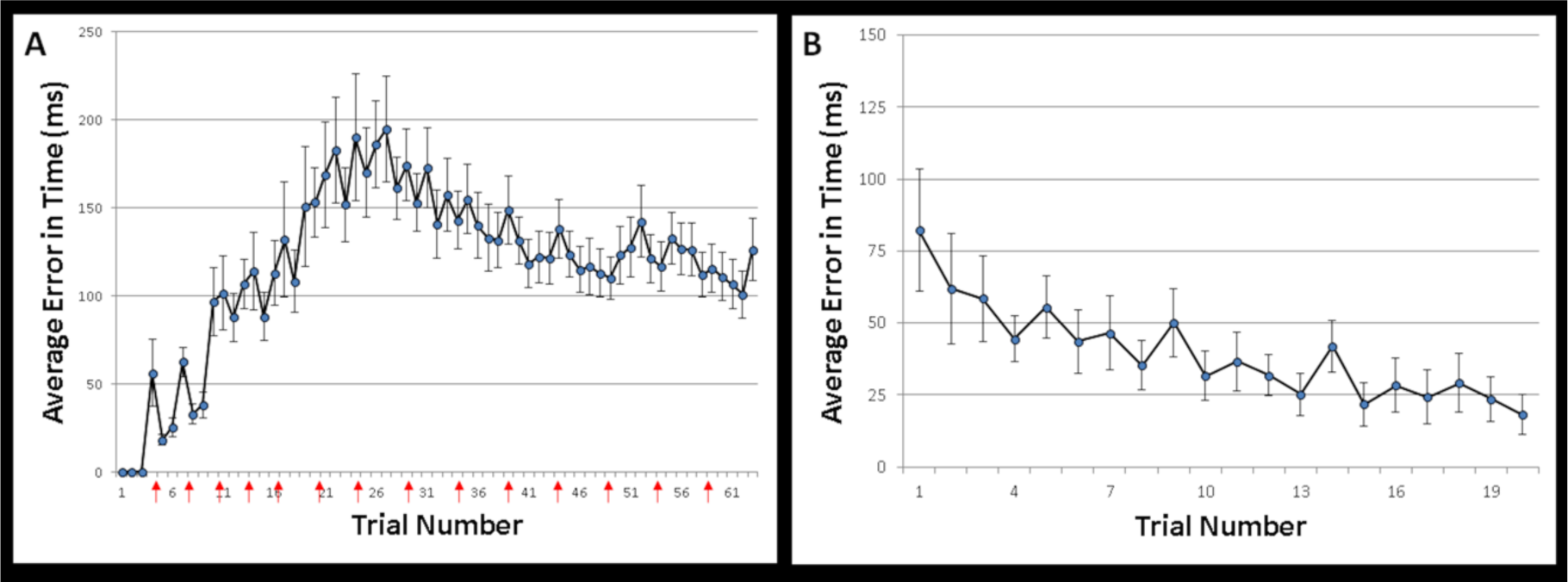
Performance in Piano Training (Timing) Accuracy in the timing of key pressing during the first (A) and second (B) learning sessions is shown. Blue circles represent the average error in time per note for each trial. (A) During the first learning session, subjects did not show a decrease in their error rate. Red arrows indicate trials in which the number of notes increased. (B) During the second learning session, subjects’ error rate decreased dramatically as they reached to an average error per note shorter than 20 milliseconds. Error bars depict standard error of mean.

#### 3.1.2 Second Learning Session

Participants that continued on to a second learning session, in which they were given feedback on the timing of key pressing, improved their timing performance dramatically: the average error per note decreased from trial to trial and by the 20^th^ trial the deviation from the correct timing decreased to less than 20 milliseconds per note (Figure 2B).

### 3.2. Accuracy Related Changes in Diffusion Properties (1^st^ session)

We performed a paired t-test on the pre- and post-learning MD images of all 32 non-musician participants in order to detect regions of learning-related MD decrease. We found a significant MD reduction in several brain regions, including the left premotor cortex and the superior part of the cerebellum, in both right and left hemispheres. In addition to evidence of structural plasticity in these motor system regions, MD reduction was also observed in the left middle temporal gyrus (Figure 3). The reduction in MD was about 2-3% in all regions (described in Figure 5). Additional smaller clusters in the right supplementary motor area and the posterior part of the right cerebellum (*P* < 0.005, cluster size > 15 voxels, Figure S1) did not exceed the corrected threshold but are shown nevertheless due to the relevance of these regions to the task.

**Figure 3:**
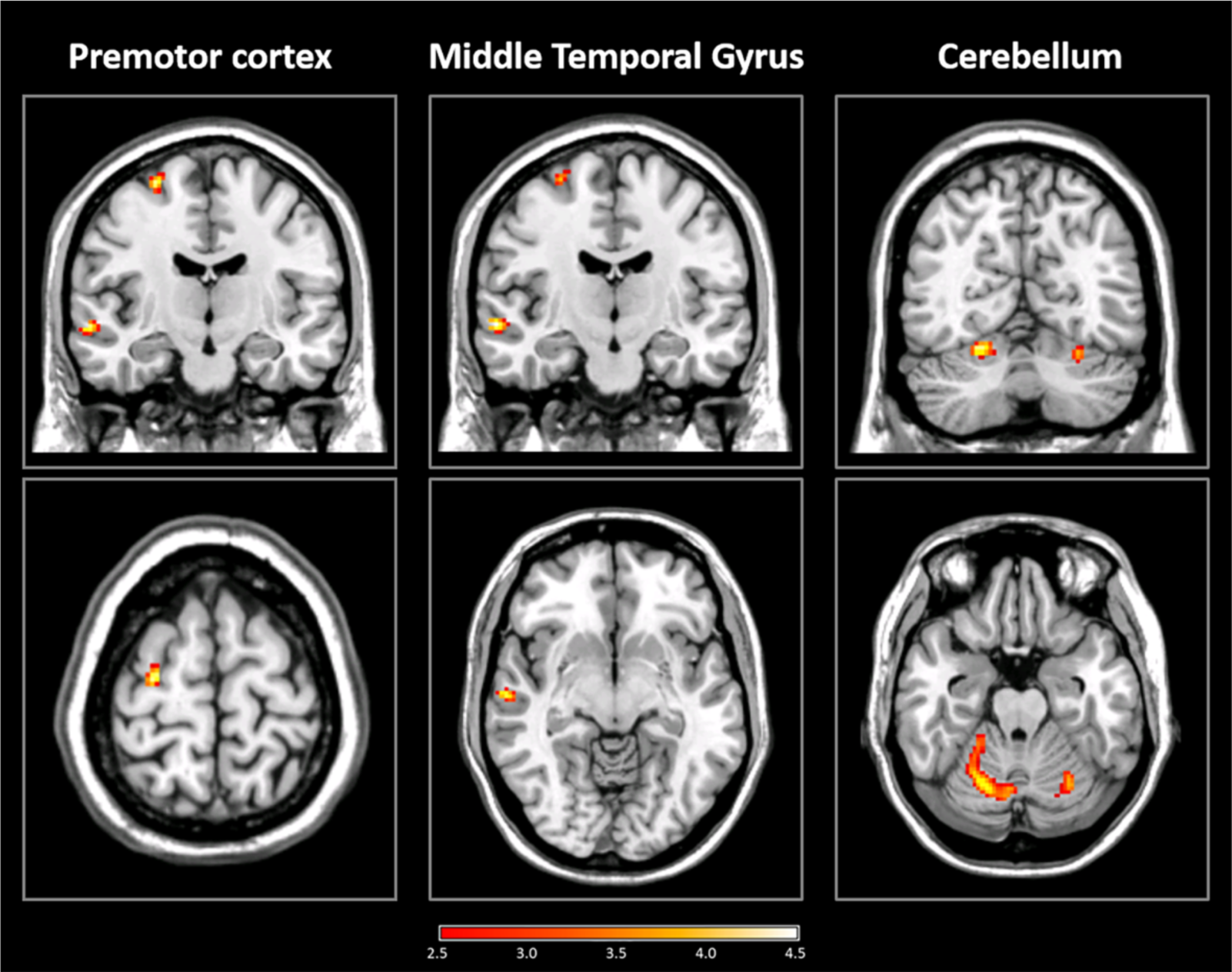
Reduction in Mean Diffusivity after Piano Accuracy Training. Structural remodeling of brain tissue, measured by DTI as a reduction in mean diffusivity (MD) after 45 minutes of training on a motor sequence learning task. A paired t-test between the MD maps before and after the learning task (first session) was performed. The statistical parametric map is presented superimposed on coronal (upper row) and axial (lower row) slices of a single-subject T1 map. Significant clusters of MD decrease were found in the left premotor cortex, left middle temporal gyrus and the cerebellum (*P* < 0.005, cluster size > 37, equivalent to *P* < 0.05, corrected for multiple comparisons). L indicates the left side of the brain; color bar represents the *T* value

**Figure 4:**
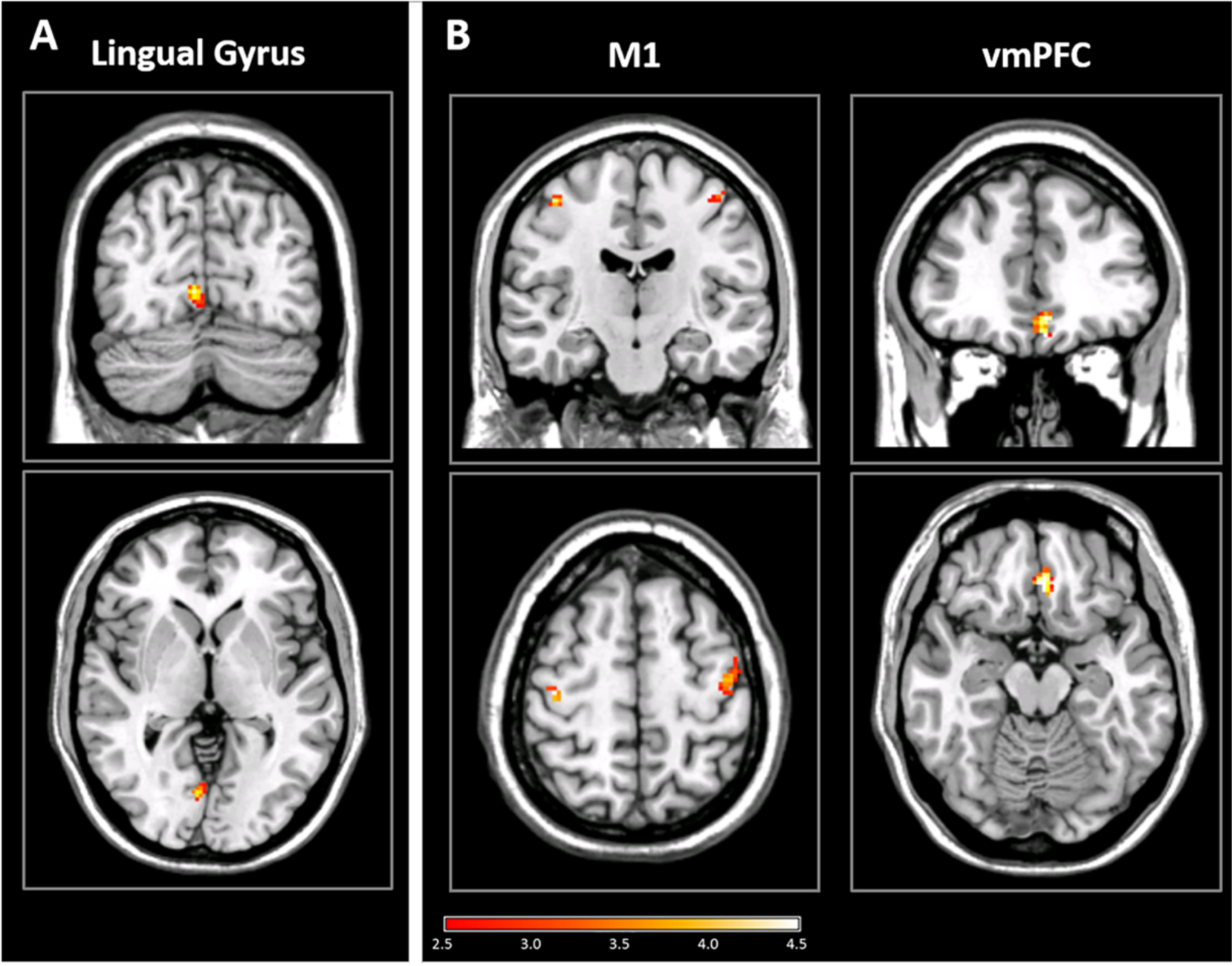
Reduction in Mean Diffusivity after Piano Timing Training in naïve participants(A) and after Training in Professional Pianists (B) Structural remodeling of brain tissue, measured by DTI as a reduction in mean diffusivity (MD) after 45 minutes of training on a motor sequence learning task. (A) A second training session in the naïve participants focused on the timing of key pressing. Analysis of Variance (ANOVA) of the MD maps before and after each learning task was performed, and post-hoc analysis revealed a significant cluster in the left lingual gyrus (*P* < 0.005, cluster size > 37, equivalent to *P* < 0.05, corrected for multiple comparisons) in which the effect was a result of a reduction in MD after the second learning session (timing). (B) Professional pianists underwent a training session similar to the first session in the naïve participants. A paired t-test between the MD maps before and after the learning task was performed. Significant clusters of MD decrease were found in the Primary motor cortex bilaterally and in the ventromedial prefrontal cortex (vmPFC). *P* < 0.005, cluster size > 37, equivalent to *P* < 0.05, corrected for multiple comparisons. The statistical parametric maps are presented superimposed on coronal (upper row) and axial (lower row) slices of a single-subject T1 map. L indicates the left side of the brain; color bar represents the statistic (*F* or *T)* value

**Figure 5:**
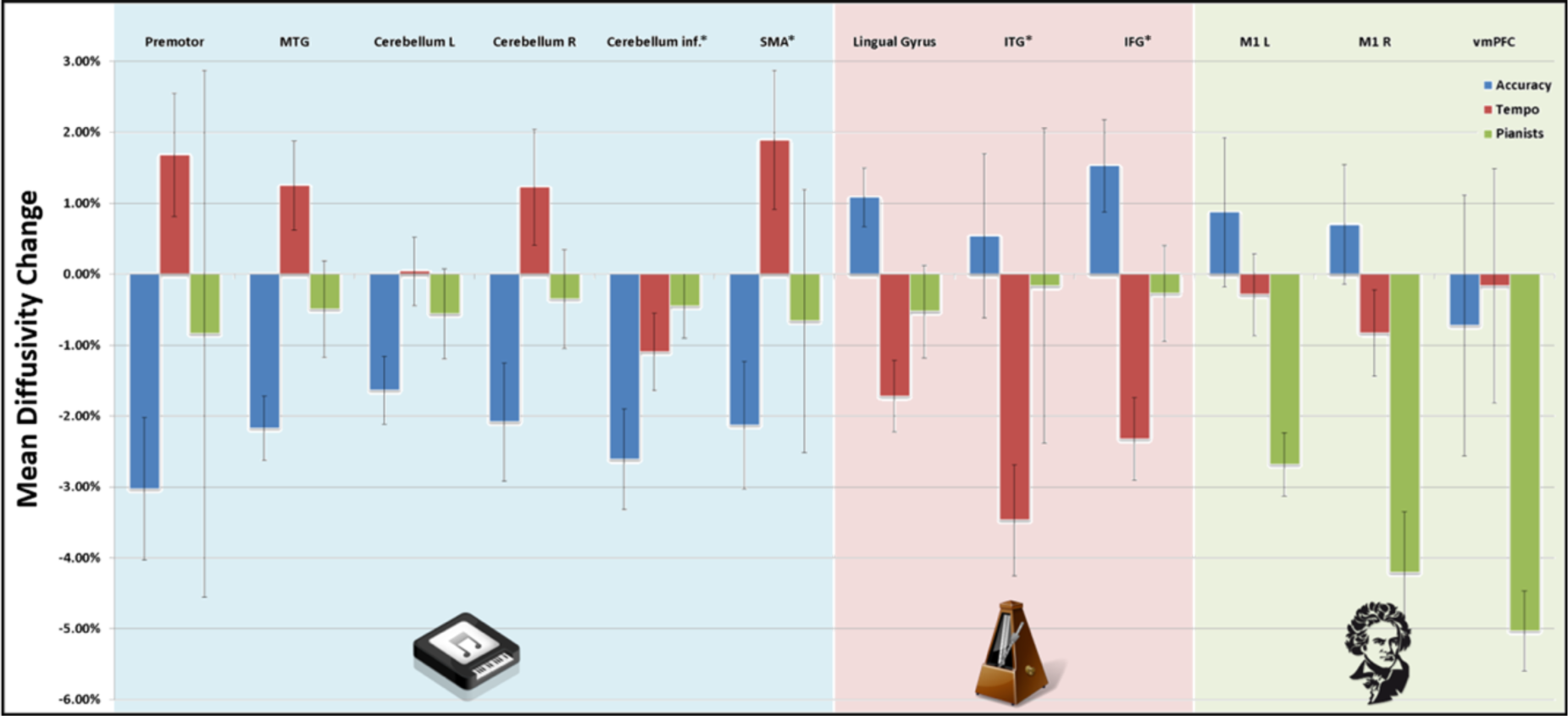
Specificity of Brain Networks Plasticity to the Learning Procedure. Significant reduction in mean diffusivity was found in different brain regions for different aspects of learning. Regions that underwent significant change after accuracy training are presented on the left, regions of significant change after rhythm/timing training are presented in the middle and regions that were found in professional pianists are presented on the right. For each of these regions, the percentage reduction in mean diffusivity ((MD after - MD before) / MD before) is shown for 3 conditions: accuracy and timing training of naïve subjects (blue and red bars, respectively) and accuracy training for professional pianists (green bar). Error bars depict the standard error of the mean.

### 3.3. Timing Related Changes in Diffusion Properties (2^nd^ Session)

We performed a repeated-measures analysis of variance (ANOVA) on the MD images of the 15 participants that performed two learning sessions and were scanned three times, in order to detect differences in structural plasticity as a result of different aspects of sequential motor learning. Post-hoc analysis revealed two types of clusters: regions that showed decreased MD in the second scan compared to the first, but less so in the third scan, indicting structural plasticity related to accuracy learning; and regions in which there was no significant change between the first and second scans yet a significant reduction in MD was observed in the third scan, indicating structural plasticity related to rhythm learning.

The comparison between the first and second scans, taken before and after the first learning session in which emphasis was put on the accuracy of key pressing, revealed evidence of structural plasticity in some of the regions that were already found in the whole cohort (see Section 3.2 above): the left premotor cortex (*F*_(2,28)_ = 7.64, *P* < 0.005), the left middle temporal gyrus (*F*_(2,28)_ = 8.68, *P* < 0.005) and the cerebellum (*F*_(2,28)_ = 7.26, *P* < 0.005). The comparison between the second and third scans, taken before and after the second learning session in which emphasis was put on the timing of key pressing, revealed evidence of timing-related structural plasticity in the left lingual gyrus (*F*_(2,28)_ = 8.87, *P* < 0.005, Figure 4A). Additional smaller clusters were revealed in the anterior part of the right inferior temporal gyrus and the left inferior frontal gyrus, but these did not exceed the corrected threshold of 37 voxels (*P* < 0.005, cluster size > 15 voxels, Figure S2).

### 3.4. Changes in Diffusion Properties in Professional Pianists

A comparison of the pre- and post-learning MD images was performed also for the professional pianists’ group. Even though they participated in the exact same task, the location of MD reduction in this group was completely different: while no change was observed in the premotor cortex or cerebellum, we detected a bilateral significant MD reduction in the primary motor cortex (M1) and the ventromedial prefrontal cortex (*P* < 0.05, corrected, see Figure 4B).

A summary of the results in all experimental groups, as well as the mean percentage change in MD values in each one of the detected brain regions, is shown in Figure 5.

## 4. Discussion

In this work we demonstrate that diffusion MRI can be used to detect rapid structural modifications in the motor system following a motor learning task. These structural modifications occur in a multi-regional fashion and at different time scales, depending on the focus of the task. Importantly, structural modifications were observed within regions of the motor system almost exclusively, even though the statistical analysis was done on the whole brain, demonstrating the specificity of the plasticity process.

First, we detected a decrease in MD in 32 non-musician participants that performed a short (~45 minutes) piano sequence learning task, focused on accuracy of key pressing. The decreased MD suggests structural tissue changes related to the learning procedure. Next, we found that a second learning session, focused on the rhythm rather than the accuracy, induced a different pattern of structural changes. Finally, we introduced the same piano learning task to professional pianists and found completely different learning-related changes in the brains of these experts compared to the non-musician participants. These results are further discussed below.

### 4.1. The Location of Brain Structural Plasticity

Most of the regions that were found to be changed are parts of the motor system, meaning that learning processes in a specific cognitive domain involve short-term structural modifications in the relevant brain regions.

The largest reduction in mean diffusivity was found in the cerebellum, as bilateral significant clusters were detected in the superior part of the cerebellar posterior hemispheres (Figure 3). A third (uncorrected for multiple comparisons) cluster was observed in inferior part of the cerebellar right hemisphere (Figure S1). The involvement of the cerebellum in motor learning is well-documented (Doyon et al., 2003; Herholz and Zatorre, 2012; Hikosaka et al., 2002; Penhune and Steele, 2012). The cerebellum, and especially its lateral parts, is considered to be essential for error-correction and error-based learning (Doya, 2000; Hikosaka et al., 2002). In a recently proposed model for motor sequence learning, the cerebellum is further suggested to be responsible for adjusting movements according to sensory inputs and for acquiring an optimal internal model for performing a sequence of movements (Penhune and Steele, 2012). Such processes are crucial for succeeding in our piano learning task. In addition, the short timescales of cerebellar modifications reported in here are also in line with previously reported evidence for rapid response of the cerebellum to explicit motor learning (Penhune and Steele, 2012), as opposed to other motor-system regions such as the putamen and primary motor cortex, that react during later stages of learning.

In addition to the cerebellum, evidence for brain plasticity was also found in the premotor cortex (Figure 3), and presumably, in a lesser extent, within the SMA (Figure S1). These regions are known to be involved in the planning of movement and were previously reported to have a role in musical training (Chen et al., 2012; Lahav et al., 2007). More specifically, the dorsal premotor cortex is considered to play a role in linking between auditory pitch information and its related key press (Chen et al., 2012; Herholz and Zatorre, 2012). The premotor cortex was found to be activated during listening to melodies that participants were previously trained to play (Lahav et al., 2007), and is considered to be involved in the integration of sensory information with motor actions (Chen et al., 2012). Taken together, our results in the cerebellum and premotor cortex demonstrate the ability of the motor system to undergo rapid structural remodeling in response to a short learning task.

Structural modifications following our motor sequence learning task were also found in the middle temporal gyrus (Figure 3), which is not a motor area per se. The temporal lobes are known to be involved in auditory performance, thus it is reasonable to assume that the middle temporal gyrus is involved in the auditory aspects of the task. The middle temporal gyrus was previously found to be sensitive to music structure (Fedorenko et al., 2012) and is considered to be part of an auditory-motor network (Bangert et al., 2006), which is required for musical abilities. In sum, learning to play music is a multisensory assignment and it is not surprising to find learning related changes in auditory regions alongside motor regions, especially regions surrounding the superior temporal sulcus which are considered to be involved in sound-action interaction (Zimmerman and Lahav, 2012).

### 4.2. Different Aspects of the Task Involve Different Brain Regions

One of the novelties of this study is the inclusion of two additional learning routines: (1) Non-musician participants underwent a second learning session, focusing on a different performance aspect - rhythm rather than accuracy. We compared the scans acquired before and after this second learning session to explore the structural plasticity that is associated with the rhythm training; (2). a preliminary cohort of professional pianists performed the first session of the piano learning task that focused on accuracy, similar to the naïve group. We compared the pre- and post-learning scans to explore the patterns of structural plasticity in the professional pianists. Figure 5 summarizes the different regions that were involved in different aspects of the task. Interestingly, the patterns of brain plasticity following rhythm training were different than those observed following accuracy training, and the plasticity patterns in the pianists’ group were different than those observed in the naïve group. These changes in the pattern of brain plasticity when the focus of learners was shifted or when learners were highly trained, highlight the dynamic nature of this phenomenon, formerly thought to require months of training in order to be detectable.

While accuracy training resulted in MD reduction mainly in motor system regions (premotor cortex and the cerebellum), rhythm training induced neuroplasticity in different locations (Figure 4A). The main change in MD was found in the lingual gyrus, a region usually associated with high-level visual processing, rather than the motor or auditory processes. The simplest explanation to this finding may involve the visual aspect of the task, in which a keyboard was presented on the screen, correct notes in a sequence were presented in blue and feedback about errors was presented to the participant in red (Corbetta et al., 1990; Tricomi et al., 2004). The visual processing of this information is relevant to accuracy training as much as for timing\rhythm training, but it is possible that the complexity of the second training might have increased the importance of the visual input and therefore the involvement of this region.

Alternatively, there is also evidence supporting the involvement of the lingual gyrus in motor (Müller et al., 2002; Parsons et al., 2005) or auditory (Bengtsson et al., 2009; Janata et al., 2002; Schmithorst and Holland, 2003) aspects of the task directly. Whether involved in the visual, motor or auditory aspects of the task, it is clear that the timing\rhythm training evoked brain networks that were less relevant to accuracy training and resulted in specific structural remodeling, as measured by a reduction in MD.

Smaller clusters of timing\rhythm training-related plasticity were found in the left inferior frontal gyrus (IFG) and the right inferior temporal gyrus (ITG) (Figure S2). While there is an extensive literature about the involvement of these regions in musical processes (Gaser and Schlaug, 2003; Penhune and Steele, 2012; Tillmann et al., 2003; Watanabe et al., 2008), conclusions about them from the current work should be done carefully as the small size of these clusters prevent them from exceeding the statistical threshold after correction for multiple comparisons. Nevertheless, the fact that only three regions showed a reduction in MD after timing training, while values in regions that were changed after accuracy training almost returned to baseline (see Figure 5), demonstrates that shifting the focus of the task to a different aspect results in a different pattern of brain plasticity.

Last, although measured in a small group of participants, results from the professional pianists provide further support for our claim that different training procedures involve different brain networks and influence brain tissue in different locations. While the professional pianists performed the same accuracy task, their cognitive requirements were totally different. They already knew the sequence and their accuracy performance showed no improvement as their level reached perfection already in the first trial and stayed the same throughout the experiment. The location of brain plasticity in these participants was very homogenous, showing no reduction in MD in any of the aforementioned brain regions but a bilateral substantial MD reduction in the primary motor cortex (M1, Figure 4B). The fundamental role of this area is to control voluntary movements, and it also has a role in motor learning (Penhune and Steele, 2012; Sanes and Donoghue, 2000). Decreased MD in the professional pianists was also found in the ventromedial prefrontal cortex (vmPFC) which might reflect changes due to automatic (Ashby et al., 2010) or habitual performance of a task (Coutureau and Killcross, 2003; de Wit et al., 2009).

### 4.3. The Temporal Dynamics of Neuroplasticity

The fact that different plasticity patterns were observed following different stages of learning reflects an important implication for the study of neuroplasticity, namely the sensitivity of diffusion MRI to dynamic, flexible brain modifications in very short time-scales. While learning-related brain changes were previously observed a week after a first learning procedure (Tavor et al., 2013), here we demonstrate a shift from an “accuracy network” to “timing network” within no more than an hour. This rapid decrease in MD values may influence the way we refer to structural brain plasticity. Obviously, interpretations of volumetric structural brain changes reported in VBM studies after months or years of training cannot explain the diffusion MRI changes reported here, and different biological substrates should be examined. The rapid modification of Astrocytes structures, described previously (Johansen-Berg et al., 2012; Sagi et al., 2012; Tavor et al., 2013), may fit the time-scale reported in the present study.

### 4.4. Technical Considerations

The current study design includes several limitations that should be noted. First, defining control conditions to the different aspects of the learning task is challenging. To overcome that, data were collected from a cohort of participants at three time-points, making it possible to analyze as an internal control condition, as performed by Thomas and colleagues (Thomas et al., 2009). Such a within-subject design with several time-points can be more powerful than a comparison between experimental groups (Thomas and Baker, 2013). Introducing the accuracy task to a small cohort of professional pianists can also be referred as a control condition and indeed analyzing this group revealed different results than the main (naïve participants) group.

In addition, the complexity of the task makes it impossible to design an equivalent animal study, as was done in a spatial learning experiment described previously (Sagi et al., 2012). This makes it harder to interpret the MRI observations and find their biological correlates. However, given the similarity of the MRI findings reported in the present study and those of Sagi et al. (i.e. MD reduction), together with the time-scale of the structural changes we observed, it is reasonable to assume that the biological substrates suggested by Sagi and colleagues (2012) are related to the MRI changes found in here as well.

Last, the statistical analysis performed in this experiment included a correction for multiple comparisons based on the combination of *P* value threshold and cluster size. The routine in which the problem of multiple comparisons should be corrected is debatable, and more strict ways to control the false-discovery-rate are sometimes expected (Bennett et al., 2009).

However, it is important to mention that the cluster size threshold used in this study was not arbitrary, but calculated specifically for our data using Monte-Carlo simulation provided by AFNI software. That way we could calculate the probability of getting a noise-only cluster and make sure the chance of it is less than 5%. The failures recently addressed by Eklund, Nichols and Knutsson (Eklund et al., 2016) are taken into consideration in this work, in aspects of cluster forming threshold and the way we estimate the smoothness of the data. Moreover, the fact that clusters were found mainly within the motor system, even though statistical analysis was performed on the entire brain, strengths our confidence that these clusters reflect true task-related brain changes rather than statistical false-positives.

## 5. Conclusions

The results reported in this work expand and elaborate our knowledge about diffusion MRI-sensitive structural brain modifications. First, we demonstrate that rapid changes in diffusion properties, indicating microstructure tissue remodeling, extend beyond traditional learning regions and occur in the neocortex and the cerebellum. Second, we show that behavioral modifications are accompanied by congruous changes to brain networks. Acquiring a new skill, specialization in a newly learned skill or practicing a well-established ability, each has its own related brain regions that undergo modifications when necessary. Last, we show that these modifications are very flexible and can be altered within a few hours. Most importantly, the current study together with previous once demonstrate the great potential of diffusion MRI for studying the dynamic nature of the adult human brain in different cognitive domains and brain systems.

## 6. Acknowledgement

The authors acknowledge with thanks the support of the Israel Science Foundation (ISF Grant no. 1314/15). The authors also thank Omri Tavor for developing the piano training software.

