## Supplementary material for "Short-Term Plasticity Following Motor Sequence Learning Revealed by Diffusion MRI": Figure S1

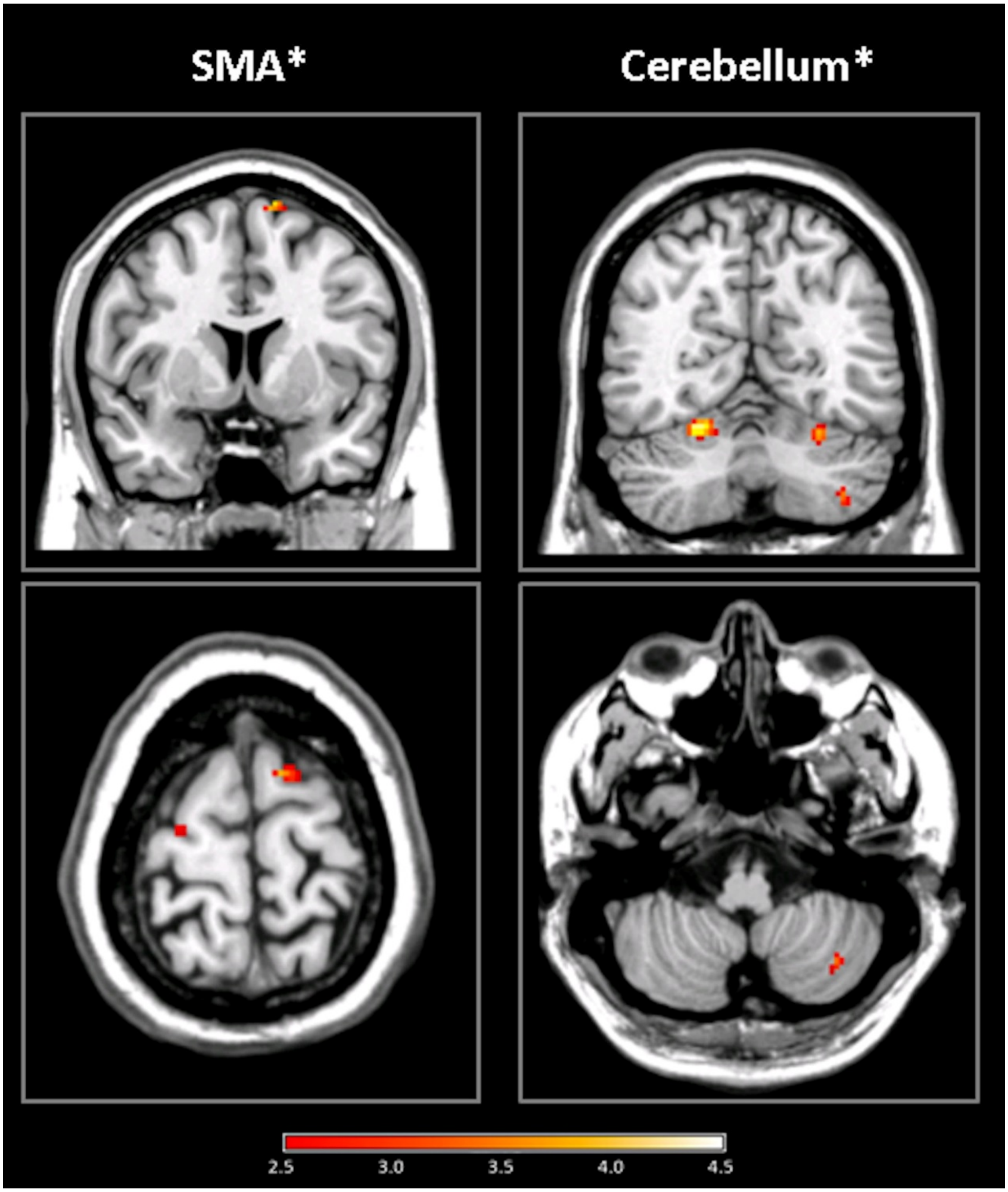


**Figure S1: Reduction in Mean Diffusivity after Piano Accuracy Training**. Structural remodeling of brain tissue, measured by DTI as a reduction in mean diffusivity (MD) after 45 minutes of training on a motor sequence learning task. A paired t-test between the MD maps before and after the learning task (first session) was performed. The statistical parametric map is presented superimposed on coronal (upper row) and axial (lower row) slices of a single-subject T1 map. Clusters found in the right supplementary motor area (SMA) and the lower part of the cerebellum ($P < 0.005$, cluster size >15) were too small to exceed a corrected threshold but are shown nevertheless due to the relevance of these regions to the task. L indicates the left side of the brain; color bars represent the *T* value
