## Supplementary material for "Short-Term Plasticity Following Motor Sequence Learning Revealed by Diffusion MRI": Figure S2

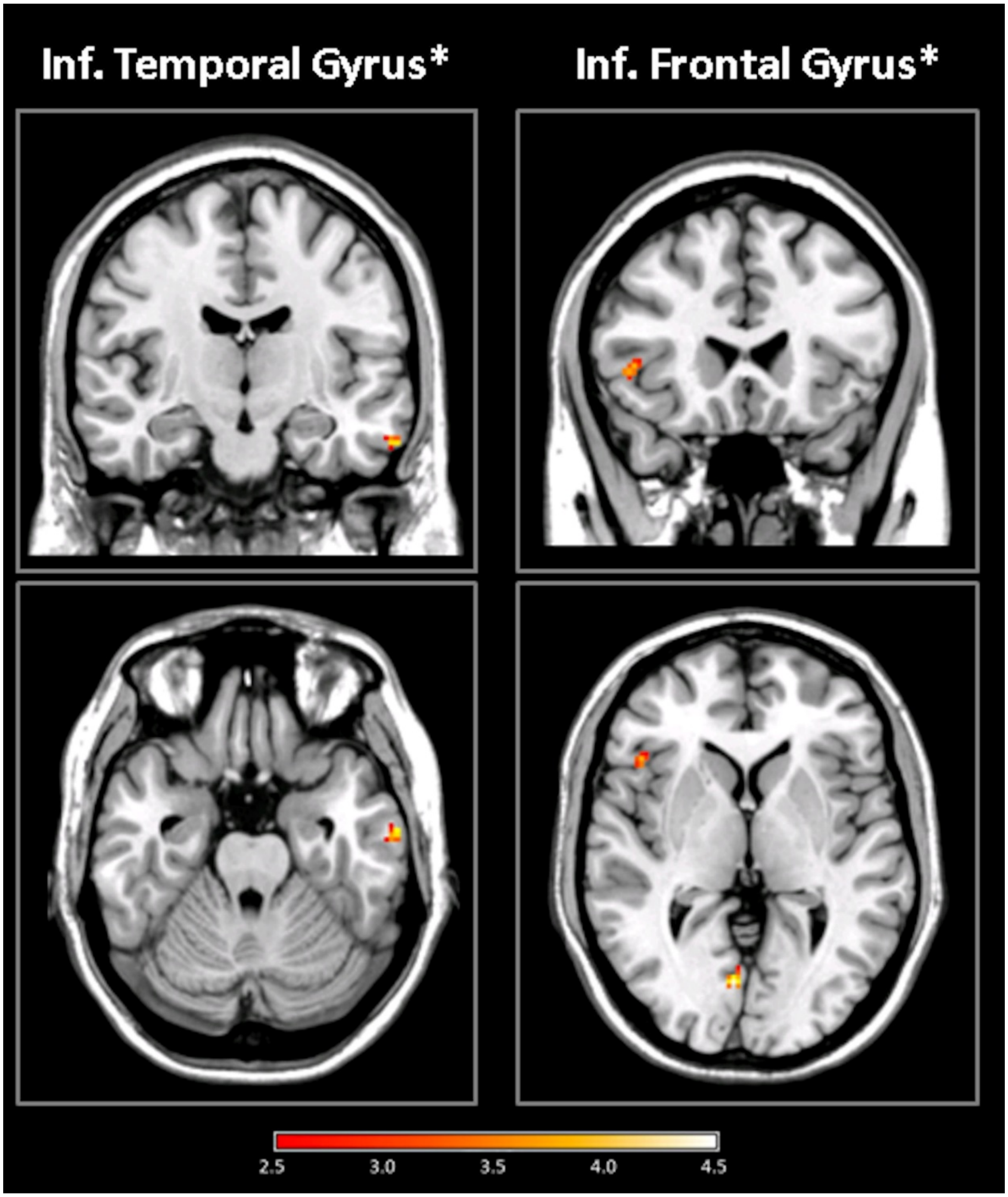


**Figure S2**: **Reduction in Mean Diffusivity after Piano Timing Training**. Structural remodeling of brain tissue, measured by DTI as a reduction in mean diffusivity (MD) after 45 minutes of training on a motor sequence learning task, focused on the timing of key pressing. Analysis of Variance (ANOVA) of the MD maps before and after each learning task was performed, and post-hoc analysis revealed clusters in the right inferior temporal gyrus and the left inferior frontal gyrus ($P < 0.005$, cluster size >15) in which the effect was a result of a reduction in MD after the second learning session (timing), yet their size was too small to exceed a corrected threshold. The statistical parametric maps are presented superimposed on coronal (upper row) and axial (lower row) slices of a single-subject T1 map. L indicates the left side of the brain; color bar represents the *F* value
